# Intrastriatal injection of amplified Lewy body dementia (LBD) fibrils in mice results in accumulation of fibrils with LBD structure and neurodegeneration

**DOI:** 10.64898/2026.10.06.757103

**Authors:** Aditi V Bagade, Rebecca Miller, Joshua T Dearborn, Katherine Basore, Brock Summers, Bradley Readnour, Matthew Koelling, Jennifer O’Shea, Elizabeth R Wright, Dhruva D Dhavale, Paul T Kotzbauer

**Author notes:** **Correspondence to:** Paul Kotzbauer, Department of Neurology, Washington University School of Medicine, Campus Box 8111, 660 South Euclid Avenue, St. Louis, Missouri 63110, USA., Dhruva Dhavale Department of Neurology, Washington University School of Medicine, Campus Box 8111, 660 South Euclid Avenue, St. Louis, Missouri 63110, USA.

## Abstract

The defining neuropathologic feature of Lewy body dementia (LBD) and Parkinson’s disease (PD) is alpha-synuclein (Asyn) fibril accumulation in Lewy bodies (LBs) and Lewy neurites (LNs). Recently, we developed a method to amplify Asyn fibrils derived from human post-mortem LBD brain tissue and determined that the structure of amplified Asyn fibrils is highly similar to the structure of fibrils directly isolated from LBD brain tissue. We examined whether injection of these fibrils into humanized Asyn mice recapitulates pathologic Asyn fibril accumulation with human LBD Asyn fibril structure. We utilized mice containing a null allele in the mouse Snca gene and a human SNCA transgene. Intrastriatal injection of amplified Asyn fibrils derived from 3 different LBD autopsy cases resulted in cytoplasmic and neuritic phospho-Asyn accumulation in the striatum, cortex, and amygdala bilaterally at 3mpi. Fibril injections produced a reduction in the number of dopaminergic neurons in the substantia nigra, with LB-like inclusions in surviving neurons, and produced impaired sensorimotor function as measured on the accelerating rotarod test. Fibril accumulation increased 3.6-fold between 1 and 3 months post-injection (mpi) and was relatively stable between 3 and 5 mpi. We amplified fibrils from mouse brain tissue 5 mpi and collected single-particle cryo-EM data. High resolution 3D structural information could not be obtained because of the lack of helical twist, but comparison of the cryo-EM 2D class averages with those derived from amplified fibrils prepared from human LBD tissue revealed multiple matching classes, indicating that the Lewy fold is maintained in the fibrils that accumulate in mouse brain tissue. Our findings demonstrate an approach to model the accumulation of Asyn fibrils with LBD structure in mice, providing a platform for investigating disease mechanisms, imaging biomarkers, and therapeutics targeting Asyn in PD/LBD.

## Introduction

Parkinson’s disease (PD) is a debilitating neurodegenerative disorder characterized by an array of clinical manifestations. It produces multiple neurologic symptoms, including movement difficulties, behavioral changes, sleep disturbances, and cognitive impairment. The motor symptoms of PD stem from progressive degeneration of dopaminergic neurons within the substantia nigra [1], while degeneration of additional subcortical and cortical neuronal populations produces nonmotor symptoms. PD is defined by the accumulation of alpha-synuclein (Asyn) fibrils in the brain, which accumulate in neuronal cytoplasmic and neuritic inclusions known as Lewy bodies (LBs) and Lewy neurites (LNs) [2] [3]. Dementia develops in up to 80% of people with PD [4] and is referred to as Lewy body dementia (LBD). It may begin concurrently with motor symptoms, as seen in Dementia with Lewy bodies (DLB), or it may develop up to 20 years after the onset of motor symptoms, as in Parkinson’s disease with dementia (PDD) [5]. Studies of genetics and disease mechanisms support a pathogenic role for Asyn fibril accumulation [6].

The development of animal models that recapitulate the accumulation of Asyn fibrils in neurons may facilitate studies of disease mechanisms and enable the evaluation of diagnostic and therapeutic strategies for PD. Since Asyn fibril accumulation is the defining pathologic feature of PD, mouse models that recapitulate the accumulation of Asyn fibrils may be particularly valuable for preclinical studies focused on Asyn as a therapeutic and diagnostic target. Intrastriatal injections of Asyn fibrils prepared *in vitro* from Asyn recombinant protein, often referred to as preformed fibrils (PFFs), can initiate cytoplasmic and neuritic accumulation of Asyn fibrils in the striatum as well as the cortex when mouse Asyn PFFs are injected into wild-type (WT) mice [7]. Fibril accumulation in this mouse model approach is likely mediated by cellular uptake of injected fibril seeds, followed by growth of fibril seeds in the presence of monomeric Asyn. Intracellular transport of fibrils may explain the accumulation of fibrils in regions distant from the site of injection. Subsequent studies have shown that PFFs prepared from human recombinant Asyn protein can also initiate fibril accumulation in both WT [7–14], and transgenic mice [15–21]. Asyn fibrils extracted from PD/LBD postmortem tissue, as well as fibrils prepared by amplifying PD/LBD fibrils can also seed Asyn accumulation in WT mice as well as mice expressing human Asyn under the Thy1 promoter and a mouse Snca gene knockout [19,22]. The morphology of Asyn inclusions resulting from the injection of extracted PD/LBD fibrils as well as amplified PD/LBD fibrils is distinct from the morphology of inclusions resulting from the injection of human Asyn PFFs [19,22].

The structure of Asyn fibrils accumulating in LBD brain tissue has been determined by cryo-electron microscopy (cryo-EM) and is distinct from the structures previously reported for Asyn fibrils produced *in vitro* under a variety of different conditions, often referred to as PFFs [23]. LBD fibril structure is also distinct from the structure of Asyn fibrils accumulating in the brain in multiple system atrophy (MSA), a neurodegenerative movement disorder defined by accumulation of Asyn fibrils in glial cytoplasmic inclusions (GCIs) observed in oligodendrocytes [24]. The overall structure of MSA fibrils differs from structures reported for PFFs, although each of the two protofilaments in MSA fibrils have a Greek key fold that is highly similar to the Greek key fold observed in many PFF preparations.

We recently reported an approach to amplify LBD fibrils, which combines fibril seeds extracted from postmortem LBD tissue with recombinant Asyn monomer and subjects the mixture to multiple cycles of sonication followed by quiescent incubation [25]. Amplified LBD Asyn fibrils comprise a combination of two-protofilament and single-protofilament fibrils. The protofilament fold of amplified LBD fibrils determined by SSNMR is highly similar to the fold of fibrils extracted from human LBD tissue [23]. Here we investigated whether amplified LBD fibrils can be utilized to produce a mouse model of Asyn fibril accumulation that recapitulates the fibril structure specific to human LBD. Based on previous work with PFFs, we utilized unilateral intrastriatal injection of fibrils. Since the primary amino acid sequence of mouse Asyn protein differs from that of human Asyn protein at 7 of 140 amino acids, we utilized a “humanized” Asyn mouse strain containing a mouse snca gene deletion and insertion of a transgene containing the human *SNCA* gene locus [26,27]. We observed progressive accumulation of Asyn fibrils following intrastriatal injection of amplified LBD fibrils, and cryo-EM 2D classification analysis indicates that LBD fibril structure is maintained during propagation of fibrils in the mouse brain. Further analysis indicates that LBD fibril accumulation produces neurodegenerative changes and behavioral impairment over 3 mpi to 6 mpi.

## Material and Methods

### Mice

PAC-Tg(SNCAWT);Snca-/- mice purchased from Jackson Laboratory were used for the experiments (Strain #: 010710) [26,27]. The Mouse Genetics Core at Washington University maintained, bred, and genotyped the mice. They were housed in a clean barrier facility with a temperature-controlled environment and a 12-hour light/dark cycle. The mice had free access to food and water and were monitored and treated at least 3 times a week for dermatitis or any other health issues. The mice with dermatitis were sprayed with Vetricyn. All animal procedures were performed in accordance with NIH guidelines [28] approved by the Institutional Animal Care and Use Committee (IACUC) at Washington University School of Medicine under the protocols 19-1098 and 23-0003.

### Recombinant Asyn monomer production

Recombinant Asyn monomer protein was produced from *E. coli* BL21 cells using the pRK172 plasmid containing the human full-length WT-Asyn construct. The detailed method to produce Asyn protein is described in [5]. Briefly, BL21 cells were transformed and grown overnight in Terrific Broth medium. After isolating bacterial pellets, Asyn was extracted using osmotic shock treatment, MgCl_2_ and streptomycin treatment, heat precipitation, and anion exchange chromatography. For use in amplified fibril production, the purified monomer was filtered through an Amicon Ultra-4, 50k (Catalog UFC805024, Millipore) cutoff filter to remove any preformed Asyn aggregates. Micro-BCA assay was used to determine the protein concentration of the Asyn monomer.

### Generation of sarkosyl-insoluble Asyn seeds

Gray matter was dissected from pathologically confirmed PD cases and extracted to generate sarkosyl-insoluble Asyn seeds for subsequent amplifications. Briefly, ∼100 mg tissue was extracted using Dounce tissue grinders (KT885300-0002) in a series of ice-cold buffers at 6 mL/g. wet wt. Concentrations: High Salt (HS) buffer (50 mM Tris-HCl pH 7.5, 750 mM NaCl, 5 mM EDTA plus 0.1% Protease Inhibitor (Sigma P2714)), HS buffer plus 1% Triton X-100, HS buffer with 1% Triton X-100 and 1 M sucrose (No Protease Inhibitor). The resulting insoluble fraction was extracted with HS buffer plus 1% sarkosyl (N-Lauroylsarcosine sodium salt, Sigma 61745) and incubated at 4 °C for 16 hours with end-over-end rotation. The resulting fraction was processed through 20 strokes of Pestle B in the Dounce tissue grinder and centrifuged at 100,000 × g at 4 °C for 20 min to isolate sarkosyl-insoluble Asyn seeds. The resulting pellet was resuspended in 20 mM HEPES, pH 7.7, + 100 mM NaCl + 0.1% Triton X-100 using the grinder at 3 mL/g wet wt. This sarkosyl insoluble fraction was subjected to a 10,000 × g spin at 4 °C for 20 min to pellet larger tissue components, and the supernatant was used for amplifications [5].

### Generation of amplified LBD fibrils using sarkosyl insoluble Asyn seeds for stereotaxic injections

The method to generate amplified LBD fibrils using sarkosyl insoluble Asyn seeds was based on methods previously described in Dhavale et al. [5], with minor modifications that included lower volumes and lower sonication intensity for each cycle. Sarkosyl insoluble Asyn seeds (5 µL) were brought up to 30 µL using 20 mM Tris HCl, pH 8.0 + 100 mM NaCl (fibril buffer). This mix was sonicated in a bath sonicator (Model Q700, Qsonica with a 5.5 inch cup horn attachment) for 1 min at amplitude 20 at 4 °C. To the sonicated seeds, 1.5 µL of 2% Triton X-100 was added to recover the sonicated tissue-derived Asyn fibril seeds. To this mixture, 50k Amicon ultrafiltered Asyn monomer was added to a final concentration of 2 mg/mL in a final volume of 100 µL. This mixture underwent quiescent incubation at 37 °C for 3 days, completing the 1^st^ cycle of sonication plus incubation. After the first cycle, the mixture was sonicated at amplitude 20 for 1 min, and an additional 100 µL of 2 mg/mL Asyn monomer was added. This mixture underwent quiescent incubation at 37 °C for 2 days (2^nd^ cycle). Then, the mixture underwent 3^rd^-cycle sonication at amplitude 20 for 1 min followed by quiescent incubation at 37 °C for 2 days, and 4^th^-cycle sonication at amplitude 20 for 1 min followed by quiescent incubation at 37 °C for 3 days. After 4 cycles, amplified LBD fibrils were stored at 4 °C if needed prior to setting up an additional three cycles of amplification to produce sufficient quantities for mouse injections. A portion of 4^th^ cycle amplified LBD fibrils (60 µL) was moved to a new tube and centrifuged at 21,000 × g for 15 min at 4 °C. The top 50 µL was removed and replaced with 90 µL of fibril buffer, and the pellet was resuspended well. The mix was sonicated (amplitude 20 for 1 min), and Asyn monomer was added to a final concentration of 2 mg/mL in a final volume of 200 µL. This mixture underwent quiescent incubation at 37 °C for 2-3 days (5^th^ cycle). It was then sonicated (amplitude 20 for 1 min), and an additional 200 µL of 2 mg/mL Asyn monomer was added, followed by quiescent incubation at 37 °C for 2-3 days (6^th^ cycle). This 400 µL 6^th^ cycle mixture was sonicated (amplitude 20 for 1 min), and an additional 200 µL of 2 mg/mL Asyn monomer was added and then incubated for 2-3 days, yielding 7^th^-cycle amplified LBD fibrils in a final volume of 600 µL. The 7^th^ cycle amplified LBD fibrils were stored at 4 °C until use. The concentration of fibrils for injections was determined by either radioligand assay or Guanidine hydrochloride (GdnHCl)-A_280_ assay, which produce similar measurements of concentration, as described previously [5].

### Cryo-EM sample preparation, data collection, and data processing

Quantifoil Copper R2/2 (Electron Microscopy Sciences) holey carbon grids were negatively glow-discharged for 60 s using a GloQube (Quorum Technologies). Between 0.1-0.5 mg/ml of amplified LBD fibrils (∼3 µL volume) in 20 mM HEPES, pH 7.7 + 100 mM NaCl + 0.008% Amphipol A8-35 were applied to the grids, and after a 20s incubation, each grid was blotted for 3 s and immediately plunged into liquid ethane using a Vitrobot Mark IV (Thermo Fisher Scientific). Grids were clipped and loaded into a 300 kV Titan Krios G3 Cryo-TEM (Thermo Fisher Scientific). Movies were recorded using a Falcon 4i Direct Electron Detector (Thermo Fisher Scientific) with a Selectris X energy filter set to a 10 eV slit width in an automated fashion using EPU software 3.11 (Thermo Fisher Scientific) with a pixel size of 0.776 Å and a nominal defocus range of −1 to −2.4 mm with a 0.2mm incremental step size. 2000 movies were acquired with a dose rate of 11.5 e-/px/s over a 2.93 s exposure time, yielding a total dose of 56 e-/Å2 over 50 fractions.

All data processing was performed in cryoSPARC [29], except for filament tracing, which was performed using crYOLO [30]. Raw movies first underwent patch motion correction and patch CTF estimation. A subset of the resulting micrographs was manually traced to generate training data for a crYOLO model. The resulting model was then used to trace filaments for the full set of micrographs, with the box distance set to 50 px. The output particle coordinates were imported into cryoSPARC, and particles were initially extracted using a 400 px box size and Fourier-cropped to 200 px. The particle stack was then subjected to 2D classification using the following custom parameters: 200 classes, maximum resolution and maximum alignment resolution of 4 Å, 2 final full iterations, 20 online-EM iterations, and a batch size of 400 per class. Following initial classification, particles were re-extracted without Fourier cropping and subjected to further 2D classification using the same custom parameters, with the initial classification uncertainty factor set to 5.

### Preparation of amplified LBD fibrils for stereotaxic injections

A sterile microcentrifuge tube was incubated with 200 µL of 2 mg/mL 50k Amicon Ultra WT Asyn monomer for 10 min to block fibril adsorption on plastic. Then, 5 µg of amplified LBD fibrils were added to the tube, and the sample was spun down at 100,000 × g for 20 min at 4 °C to pellet the amplified LBD fibrils. The supernatant was removed, and the pellet was washed with 150 µL of sterile Phosphate-Buffered Saline (PBS) (14190-136, Gibco). The sample was then spun down again, resuspended in sterile PBS (minimum volume 30 µL), and sonicated for 2 min at amplitude 50 in a sonicator with cup horn attachment (Q700, QSonica). Sonicated fibrils were pulse centrifuged in a benchtop centrifuge to collect any droplets formed at the top of the tube during sonication, and a sample was taken for stereotaxic injections.

### Stereotaxic injections

Stereotaxic surgery was performed according to Zhang et al [11]. Mice at 7-8 months were anesthetized with isoflurane and administered Buprenorphine-ER 0.1 mg/kg by intraperitoneal injection for pain relief. We performed unilateral stereotaxic injections on the left side with 5 μg of amplified LBD fibrils.

Animals receiving sterile PBS were treated as vehicle control. Injections were done at the coordinates −2 mm media/lateral, 0.2 mm anterior/posterior, −3.2 mm dorsal/ventral relative to Bregma. The fibrils were injected at the rate of 1 μL/min with a total volume of 2.5 μL. Mice were monitored daily until 10 days post-surgery and then were observed three times per week until sacrificed.

### Brain tissue collection and sectioning

Mice were sacrificed at 7 days, 1, 3, 5, 7, and 14 mpi. Mice were euthanized by decapitation under anesthesia with isoflurane and the brains were collected and rinsed in ice-cold PBS. Coronal sections were cut at Bregma, ± 3.0 mm for pSyn ELISA, pSyn stain, and tyrosine hydroxylase (TH) stain. These sections were flash frozen in a dry-ice 2-methylbutane bath (Sigma 270342). These sections were then embedded in Tissue-Tek (OCT compound, Sysmex Digitana) and snap-frozen in a dry-ice/2-methylbutane bath. The OCT blocks were cryostat sectioned at 16 μm, air-dried for 15 min, and stored at −80 °C until further use.

### Immunofluorescence staining

The staining was done by fixing the fresh frozen sections in 0.1% paraformaldehyde in PBS for 15 min, followed by quenching autofluorescence with 1X Trueblack (Biotium 23007) for 15 min, and then incubated in 70% EtOH for 20 min. The slides were incubated with primary antibodies: biotin antiphospho-α-syn (S129) antibody; anti-tyrosine hydroxylase (TH) for 16 h at 4 °C **(Table S2).** The secondary antibodies were incubated for 1 h at RT **(Table S2)**. The sections were mounted with in situ mounting media with DAPI Fluoromount G (Southern Biotech 0100-20) and coverslips for microscopy. Images were captured with a Zeiss Axioscan 7 microscope at 10× magnification and Nikon TE 2000-U at 40X.

### ELISA

pSyn ELISA: Cortex and striatum were dissected out from the frozen OCT block using a dissecting stereomicroscope (Olympus SZX-ILL K1000, V lux 1000). Sequential extractions with high salt, Triton X, sucrose, and RIPA were performed to isolate insoluble fractions. These fractions were solubilized in 2% SDS with sonication and used for ELISA as described earlier [31].

### Stereology

Mouse brains 7 mpi were sectioned, containing the substantia nigra pars compacta (SNpc) −2.69 mm to −4.03 mm from Bregma (1:3 series). We performed unbiased stereology with a Nikon Eclipse E800 microscope having Stereoinvestigator (MBF Bioscience) optical fractionator probe [32]. A blind study was performed. We first drew contours around SNpc at 4X objective[33], then selected a grid size of 100 μm×100 μm. This grid was placed over the drawn contour and counting boxes of 50 μm×50 μm were randomly selected by the program to quantify SNpc. We used a guard zone of 1.5 μm for each top and bottom, and an optical dissector height of 20 μm. The measured section thickness was approximately 22 μm. TH neurons within the counting frame were counted while focusing through a z-stack. Counts were performed using a 60X oil immersion objective and the optical fractionator to estimate total stained neurons. Variability was assessed with the Gundersen coefficient of error (≤0.1). One section from one mouse brain was lost due to breakage during focusing, which was likely caused by oil-induced misalignment of the slide, and was excluded from analyses.

### Behavior testing

For all behavior testing, we injected mice with Human-amplified LBD 1 sarkosyl fibrils, referred to as fibrils in the figures, and injected Phosphate Buffer Saline (PBS) as a vehicle control. For 3 mpi fibrils injected mice: 7 females, 8 males; Control: 9 females, 6 males. For 6 mpi, fibrils injected mice: 8 males, 7 females; control: 8 females and 5 males.

Accelerating Rotarod. At both 3 mpi and 6 mpi, all mice were tested for motor coordination and balance using an accelerating rotarod apparatus (AccuRotor EzRod, Omnitech Electronics Inc., Columbus, OH) and associated software (Fusion v6.5 r1200, AccuRotor Edition, Omnitech Electronics Inc., Columbus, OH). The apparatus consists of a rotating rod, 3 cm in diameter and etched with grooves, elevated at a height of 36 cm. Briefly, mice were tested for two consecutive days (3 trials/day, 30-minute inter-trial interval) at a speed accelerating from 4 to 40 rpm over 3 minutes. Mice were allowed to stay on the rotating rod until they fell off, or the timer reached 180s. Data (seconds spent on the rod) were analyzed via repeated-measures ANOVA, comparing treatment groups across repeated trials at each timepoint.

Locomotor Activity. To evaluate levels of locomotor activity at the earlier and later timepoints, mice were observed across 1 h freely exploring a rectangular transparent polystyrene enclosure measuring 47.6 cm long, 25.4 cm high, and 20.6 cm tall. As described previously [34], each cage was surrounded by two frames of photocells that created a lower and higher 4 x 8 matrix of photobeams. Data were collected as photobeam breaks via MotorMonitor software (Hamilton-Kinder LLC, Poway, CA) and converted to distance traveled (in cm) as well as time spent (in sec) in either a central zone measuring 33 x 11 cm or an edge/peripheral zone that was 5.5 cm wide along the outermost edge of the enclosure. Variables analyzed included total distance traveled, time spent in each zone, and rearing activity. Groups were compared via student’s t-test with significance indicated by a p value <.05.

Sensorimotor Battery. Balance, limb coordination, and strength were evaluated in mice over a brief battery of tests as described elsewhere [35]. At both timepoints, mice were assessed for pole turnaround, platform balance, pole climbdown, and walk initiation. Each assay was administered twice, and the data are reported as the mean of the two trials. When data was distributed normally, groups were compared via t-test. When assumptions of normality were not met, differences were measured using a Mann-Whitney U test.

Digigait. Gait performance was measured as detailed previously [36]. Using the Digigait Gait Analysis System (Mouse Specifics Inc.), gait metrics were collected at 3 and 6 mpi timepoints and at two speeds: 30cm/s and 40cm/s. Briefly, mice were first habituated to the apparatus on the stationary treadmill for 7 minutes, then gradually the treadmill was sped up until the mouse could sustain a run at 20cm/s. For mice that reached this threshold, the speed gradually increased to 30cm/s at which point video was recorded for analysis. On the following day, this process was repeated, with the speed gradually increasing to 40cm/s. A successful run consisted of approximately 8-10 consecutive strides in a relatively straight line, and the aforementioned software analyzed gait measures including stance width, stride length, paw angle, and others for both forelimbs and hindlimbs. Data was not scaled to body length, as the mean did not differ between groups at 40cm/s, 3mpi (Control 8.14 ± 0.33 vs fibrils 8.12 ± 0.32) or at 6mpi (Control 8.41 ± 0.36 vs fibrils 8.49 ± 0.36).

### Extraction of Insoluble Asyn fibrils from mouse cortex

Cortex was dissected from 5 mpi mice injected with amplified LBD fibrils generated using human LBD sarkosyl insoluble Asyn seeds. About 100 mg of cortex was dissected from multiple injected mice and pooled. Dissected gray matter was extracted using a Dounce Homogenizer in a series of buffers (HS buffer, HS buffer plus 1% Triton X-100, HS buffer with 1% Triton X-100 and 1 M sucrose) to generate insoluble fraction Asyn seeds from fibril-injected mice.

### Amplification of fibrils from insoluble fraction Asyn seeds from fibril-injected mice

Asyn seeds isolated from mouse cortex were amplified using purified Asyn monomer using the method described in Dhavale et al. [5]. Briefly, 10 µL of insoluble Asyn seeds were brought to 30 µL in 20 mM Tris-HCl, pH 8.0 plus 100 mM NaCl buffer (fibril buffer) and sonicated for 2 min at amplitude 50 in a bath sonicator with a cup horn attachment (Q700, Qsonica). After addition of 1.5 µL of 2% Triton X-100, 50k Amicon ultrafiltered Asyn monomer was added to a final concentration of 2 mg/mL in a final volume of 100 µL. This mix underwent quiescent incubation at 37 °C for 3 days (1^st^ cycle). At the end of 3 days, the sample was sonicated (1 min at amplitude 50), 300 µL of 2 mg/ml monomer was added, and the mix underwent quiescent incubation at 37 °C for 2 days (2^nd^ cycle). Sonication and incubation cycles were repeated for cycles 3 and 4. At the end of the 4^th^ cycle, the sample was stored at 4 °C. Further expansion to 6 cycles was performed by combining 60 µL of the 4^th^ cycle with 40 µL of fibril buffer and sonicating the mixture for 1 min at amplitude 20. To this mixture, Asyn monomer was added to a final concentration of 2 mg/ml in a final volume of 400 µL in fibril buffer. This mix was quiescently incubated at 37 °C for 2 days (5^th^ cycle). The sample was then sonicated for 1 min at amplified 20 and additional Asyn monomer and fibril buffer were added to bring the final concentration of Asyn monomer to 2.5 mg/ml in a total volume of 800 µL. This mixture was incubated at 37 °C for 2 days to complete 6 cycles of incubation. After 6 cycles, fibrils amplified from mouse brain cortex were stored at 4 °C until used for single particle cryo-EM.

## Results

### Production of amplified LBD fibrils from human LBD postmortem tissue

We utilized our previously reported amplification protocol, with a few minor modifications, to produce amplified LBD fibrils for intrastriatal injections in mice. To isolate *ex vivo* fibril seeds for amplification, we prepared sarkosyl-insoluble fractions from frozen postmortem brain tissue by performing an additional extraction in 1% sarkosyl following the preparation of the insoluble fraction used in the previously reported protocol [5]. The rationale for this additional extraction step was to further reduce the amount of human brain-derived material present in the amplified LBD fibril preparations used for injections.

We used single-particle cryo-EM analysis to confirm that amplified LBD fibrils produced from sarkosyl-insoluble fractions were structurally similar to those produced from the previously reported [5] standard insoluble fractions **(Fig. S1)**. We were unable to obtain high resolution 3D density maps due to the lack of helical twist for amplified LBD fibrils. Instead, we compared 2D class averages for amplified fibrils produced with standard insoluble fractions from the human LBD1 case that was previously used for SSNMR structural characterization [5] **(Fig. S2a)** to those produced with sarkosyl-insoluble fractions from the LBD1 case and two additional LBD cases (LBD7, LBD8) **(Table S1, Figs. S2b - d)**. All of these amplified fibril preparations comprise a mix of single and two-protofilament fibrils, with no helical twist evident in any of the class averages. The structural similarity of the amplified fibrils prepared from the LBD1 insoluble fraction and from the three sarkosyl-insoluble fractions is supported by the identification of 2D class averages with highly similar density patterns, which likely represent similar views of fibrils with the same protofilament fold seen in human fibrils [5,23].

### Time course and pattern of α-Synuclein accumulation in mice following stereotaxic injection of amplified LBD fibrils

We performed unilateral stereotaxic injection of amplified LBD fibrils into the left striatum of PAC-Tg(SNCAWT);Snca-/- mice [26,27]. It contains a P1 artificial chromosome transgene containing the human WT Asyn gene locus and promoter region. In addition, it has a null mutation in the mouse Asyn gene (Snca), eliminating protein expression.

Anti-pAsyn immunostaining was observed near the injection site at 1-week post-injection **(Fig. S3)**. The density of pAsyn immunostaining progressively increased at 1 and 3mpi and was relatively stable between 3 and 5 mpi. Pathologic Asyn accumulation, which included filamentous strands accumulating in the cytoplasm of neurons as well as neuritic accumulation, was observed in the striatum, cortical regions from the retrosplenial to the ectorhinal area, cortical amygdalar area, and amygdala bilaterally, with a higher density of accumulation ipsilateral to the injection site (**Fig. 1a, b)**. We observed similar Asyn fibril accumulation in mouse brain following injection of amplified fibrils prepared from three different human LBD autopsy cases **(Fig. S4)**.

**Figure 1:**
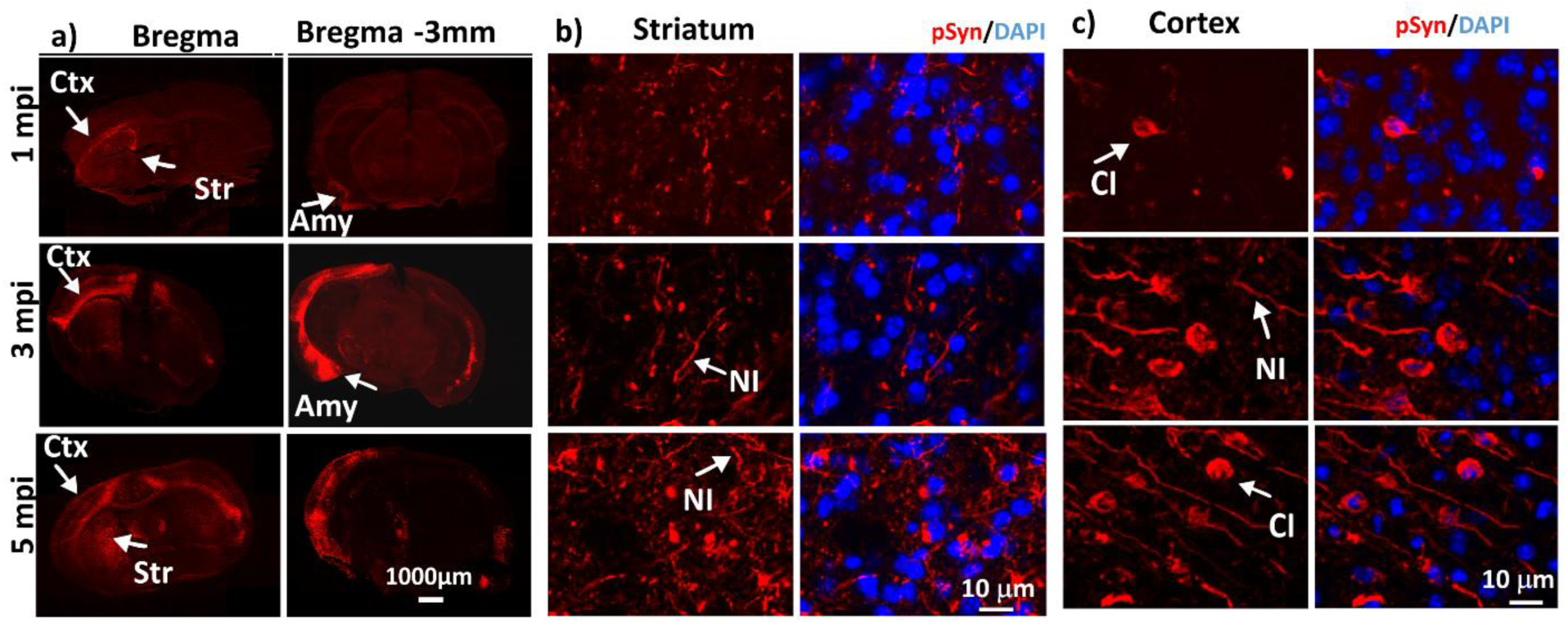
Intrastriatal injection of amplified LBD Asyn fibrils initiates neuritic and cytoplasmic accumulation of Asyn in PAC-Tg(SNCAWT);Snca-/- mice: Low-magnification images (10X objective) showing immunostaining with anti-pAsyn antibody at 1, 3, and 5 mpi of fibrils (a). Higher magnification images (40X objective) from striatum (b) and cortex (c) following pAsyn immunostaining and DAPI labeling of nuclei. Abbreviations: Cortex (Ctx), Striatum (Str), Amygdala (Amy), Cytoplasmic Inclusions (CI), Neuritic inclusions (NI); pAsyn (Red); DAPI (blue). Scale bars: 1000 μm (a), 10 μm (b), 10 μm (c). Similar anti-pAsyn immunostaining results were observed for n=20 mice at 3mpi, n=15 at 1mpi and n=13 at 5mpi.

Asyn fibril accumulation, as assessed by pAsyn immunostaining, persisted for at least 14 mpi, the latest point examined. Pathologic pAsyn accumulation was not observed in mice injected with PBS control between 1 week and 14 mpi **(Fig. S5)**. The morphology of pAsyn immunostaining evolved with respect to time post-injection. At 1 mpi, short neuritic processes were observed in the striatum and cortex, with additional infrequent filamentous strands in neuronal cytoplasm. At 3 mpi, pAsyn accumulated in frequent long neuritic processes in the cortex as well as filamentous strands in the cytoplasm of neurons, while the striatum showed less frequent neuritic accumulation (**Fig. 1**). Persistence of Asyn fibril accumulation was observed at 14 mpi, with cytoplasmic accumulation that appeared more consolidated into round inclusions **(Fig. S5)**.

We utilized a sandwich ELISA for pAsyn to measure insoluble pAsyn fibril accumulation in cortex and striatum. Levels of insoluble pAsyn were higher in cortex compared to striatum and higher on the ipsilateral side compared to the contralateral side. Within the ipsilateral striatum and cortex, a 3.6-fold increase in median pAsyn levels was observed between 1 and 3 mpi. It was relatively stable between 3 and 5 mpi. The median level of pAsyn was 1.8 times higher in cortex than striatum at 5 mpi on the ipsilateral side (**Fig. 2**). Insoluble pAsyn levels in mouse tissue were similar to levels observed for human LBD tissue (0-500ng/g wet wt.) [31].

**Figure 2:**
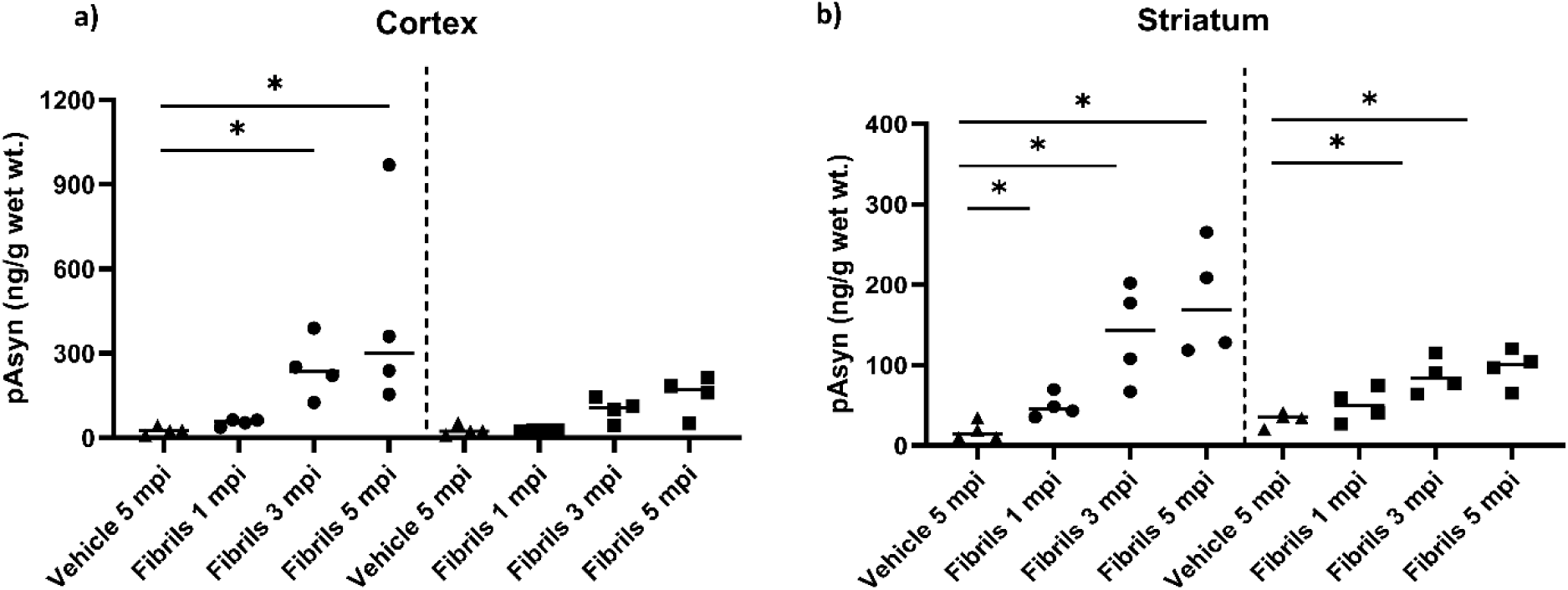
Quantification of insoluble pAsyn accumulation with a sandwich ELISA to define spatial and temporal patterns of LBD Asyn fibril accumulation following unilateral intrastriatal injection of amplified LBD Asyn fibrils. Analysis for cortex at 1, 3, and 5 mpi (a). Analysis for striatum at 1, 3, and 5 mpi (b). Samples below the LLQ were assigned half the value of the LLQ, which was 9.99 ng/g. Data to the left of the dotted line correspond to the side ipsilateral to the injection, whereas data to the right correspond to the contralateral side. The median value is shown as a horizontal bar within each group.

### Pathologic Asyn accumulation and loss of dopaminergic neurons in the substantia nigra

pAsyn immunostaining revealed neuritic and neuronal cytoplasmic accumulation in the ipsilateral substantia nigra at 3 and 5 mpi. pAsyn accumulation was also observed in the contralateral substantia nigra at 5 mpi (**Fig. 3a-d)**.

**Figure 3:**
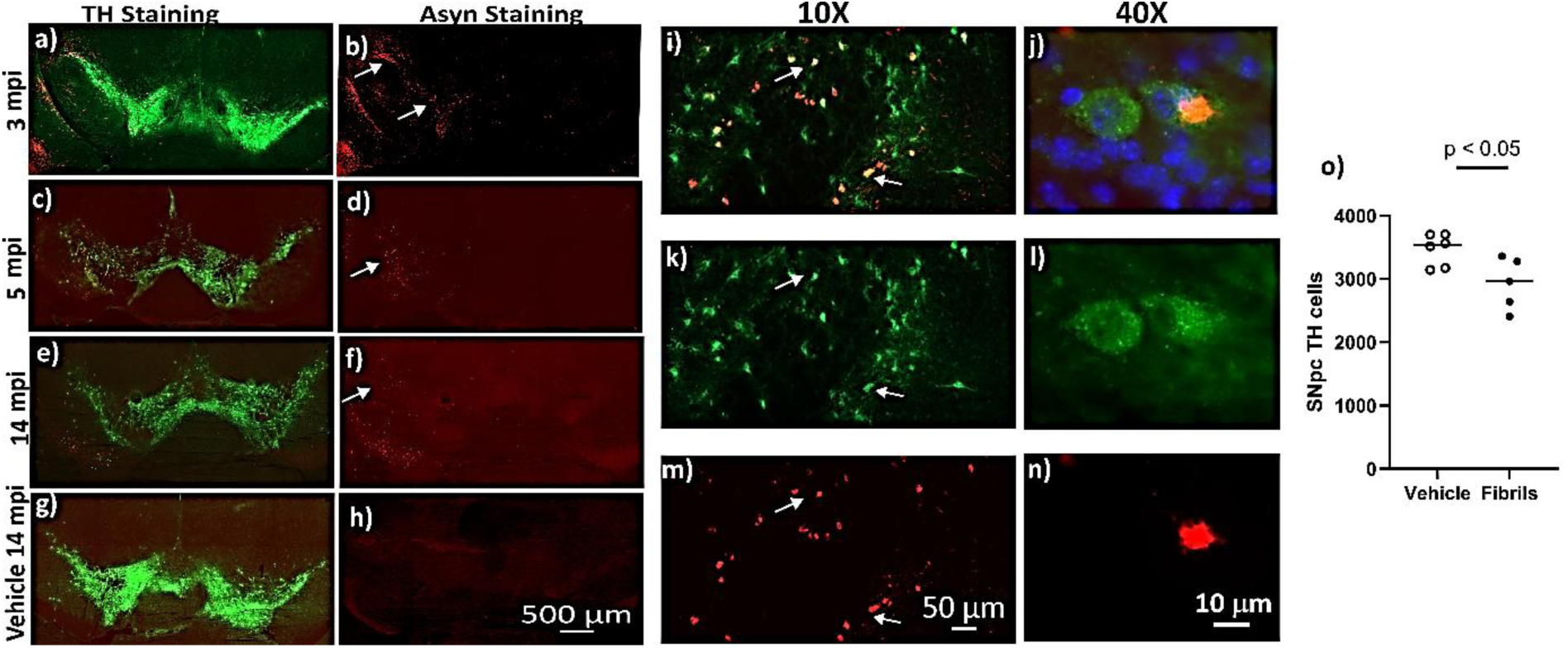
Accumulation of pAsyn in LB-like cytoplasmic inclusions in substantia nigra dopaminergic neurons on the ipsilateral side post injections. Leftmost panel: Representative images of anti-TH stained dopaminergic neurons (green; a, c, e, g) and anti-pAsyn staining (red with white arrows; b, d, f, h). Middle panel (i-n): Colocalization of dopaminergic neurons (green), pAsyn (red), and DAPI nuclear stain (blue) (i,j); dopaminergic neuron staining (k,l); pAsyn immunostaining (m,n) at 5mpi. The right panel (o) shows stereological estimates of TH-immunoreactive neurons in SNpc at 7 mpi, which shows a significant reduction in the number of dopaminergic neurons (unpaired t-test p= 0.0246, t=2.695, df=9). Scale bars: 500 μm (a-h), 50 μm (i, k, m), 10 μm (j, l, n).

Double-label immunofluorescence staining of tyrosine hydroxylase (TH), a marker for dopaminergic neurons, and pAsyn demonstrated pAsyn accumulation in dopaminergic neurons in the substantia nigra pars compacta (SNpc) at 3 through 14 mpi. (**Fig. 3a-f**). The mice injected with PBS control did not show any pAsyn accumulation up to 14 mpi (**Fig. 3 g, h)**. High-magnification images further showed colocalization of TH+ dopaminergic neurons with pAsyn immunostaining (**Fig. 3 i - n)**. We utilized stereology to quantify the number of SNpc dopaminergic neurons at 7 mpi and observed a significantly lower number of SNpc TH neurons (**Fig. 3 O)** on the ipsilateral side of LBD fibril-injected mice compared to vehicle controls (p= 0.0246).

### Sensorimotor deficits resulting from injection of amplified LBD fibrils

We used a panel of behavior tests to evaluate whether pAsyn fibril accumulation was accompanied by neurological impairment. Mice injected with LBD fibrils produced from sarkosyl-insoluble fractions had impaired performance compared to vehicle control mice on the accelerating rotarod. Both groups of mice showed improved performance with repeated trials, but fibril-injected mice were unable to stay on the rotating rod for as long as control mice at 3 mpi [F(1,28) = 5.97, p = 0.021]. A similar difference in time on the rotating rod was observed for the two groups at 6 mpi, but the difference was not significant [F(1,26) = 2.97, p = 0.096] (**Fig. 4 a, b)**. There was no statistical effect of sex on rotarod performance at either timepoint.

**Figure 4:**
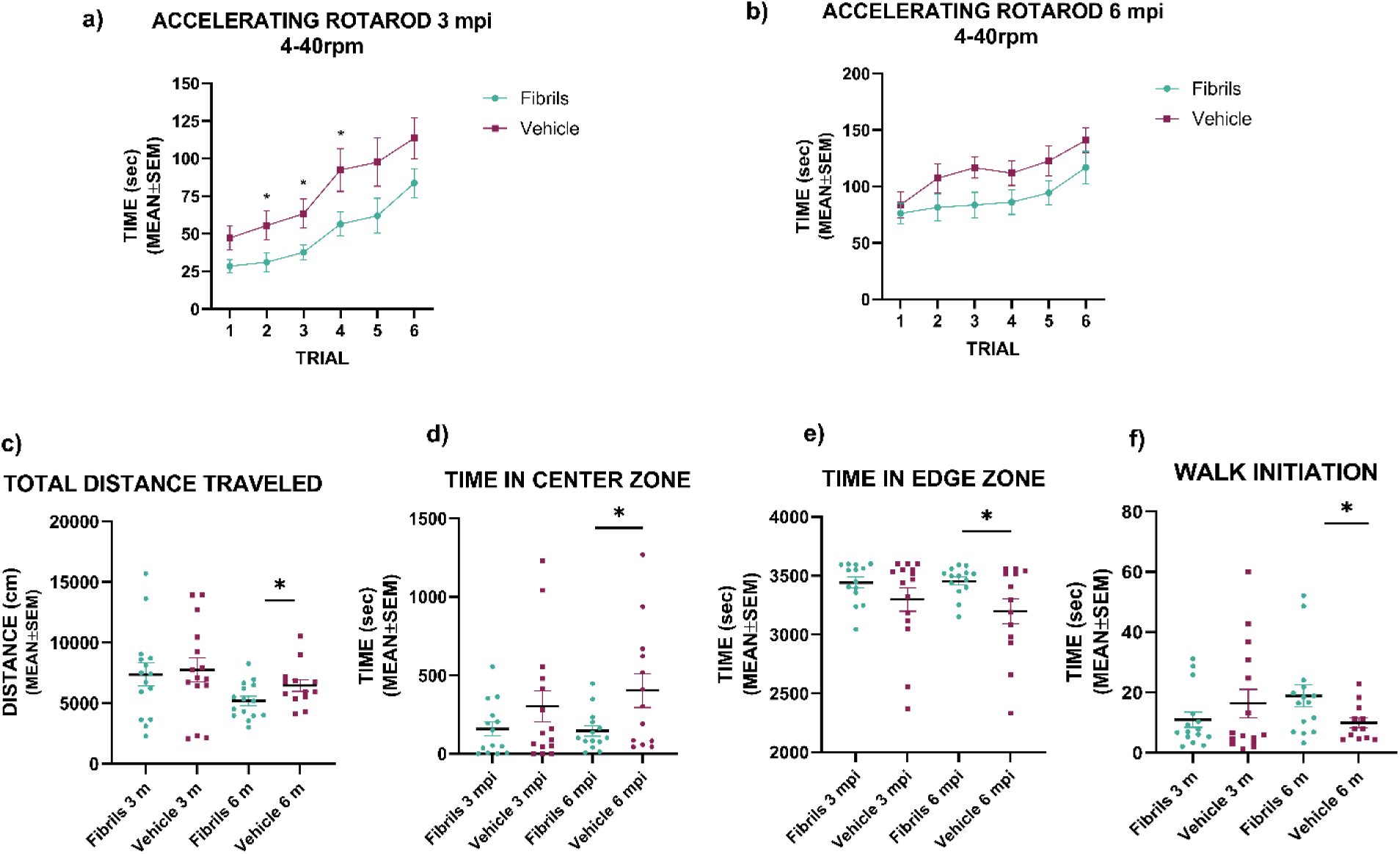
Impaired performance on the rotarod test and altered spontaneous activity patterns following intrastriatal injections of amplified LBD Asyn fibrils. At 3 mpi, mice injected with amplified LBD fibrils had impaired performance on the accelerating rotarod, as compared to mice injected with control [F(1,28) = 5.97, p = 0.021] (a). Similar differences were observed for the two groups at 6 mpi (b), but the difference was not significant [F(1,26) = 2.97, p = 0.096]. Measures of exploratory locomotor activity were similar for fibril-injected and control mice at 3 mpi, but fibril-injected mice had lower levels of activity as measured by distance traveled at 6 mpi (c-f). They also spent less time in the center zone and more time in the edge zone and were slower to initiate walking in an open field (c-f). Statistical test: Repeated measures ANOVA for (a) and (b); Independent samples t-test at each time point for (c-e); Mann-Whitney test for (f).

Fibril injections also produced differences in spontaneous locomotor activity. We assessed spontaneous locomotor activity over 1 hr at 3 and 6 mpi. There were no differences between groups at 3 mpi. However, at 6 mpi, fibrils-injected mice traveled significantly shorter distances compared to control mice (t = 2.07, df = 26, p = 0.048). In addition, fibrils-injected mice spent more time in the edge zone, and less time in the center zone, compared to control mice (t = 2.35, df = 25, p = 0.027 for both) (**Fig 4**). Sex did not significantly affect the locomotor activity of fibril-injected mice. Fibril-injected mice also took longer to initiate movement at 6 mpi (Mann-Whitney *U* = 49, *p* = 0.046) but not at 3 mpi, with no relationship to sex. Sensorimotor tests assessing pole turnaround, platform, pole climbdown and rearing time showed a trend where mice injected with fibrils performed worse than the control mice injected with PBS. Gait analysis showed no differences in stride frequency, length and swing/stand times following fibril injection, but did show significant differences at 3 mpi for the paw angle and stance width (**Fig. S6).**

### The structure of fibrils that accumulate in mouse brain tissue is highly similar to the structure of amplified LBD fibrils

To evaluate the structure of fibrils that accumulate following intrastriatal injection of amplified LBD fibrils, we extracted mouse brain tissue at 5 mpi and used the insoluble fraction from mouse brain tissue to amplify Asyn fibrils. We then analyzed the amplified fibrils with single-particle cryo-EM to evaluate their structure. 2D classification analysis indicated that all fibrils had little to no twist, as we previously observed for amplified LBD fibrils and as has also been observed for the majority of fibrils extracted from LBD brain tissue [5,37]. Due to the lack of helical twist, we were not able to obtain high resolution 3D density maps from the cryo-EM data, but comparison of the 2D class averages of the fibrils amplified from mouse brain tissue **(Fig. S7)** to those amplified from human LBD brain tissue **(Fig. S2a)** revealed multiple matching classes, consistent with the preservation of the LBD fibril fold (**Fig. 5**). Although the lack of twist currently limits 3D helical reconstruction by single-particle cryo-EM, this comparison of 2D class averages is consistent with maintenance of the LBD fold in fibrils that accumulate in mouse brain tissue following injection of amplified LBD fibrils.

**Figure 5:**
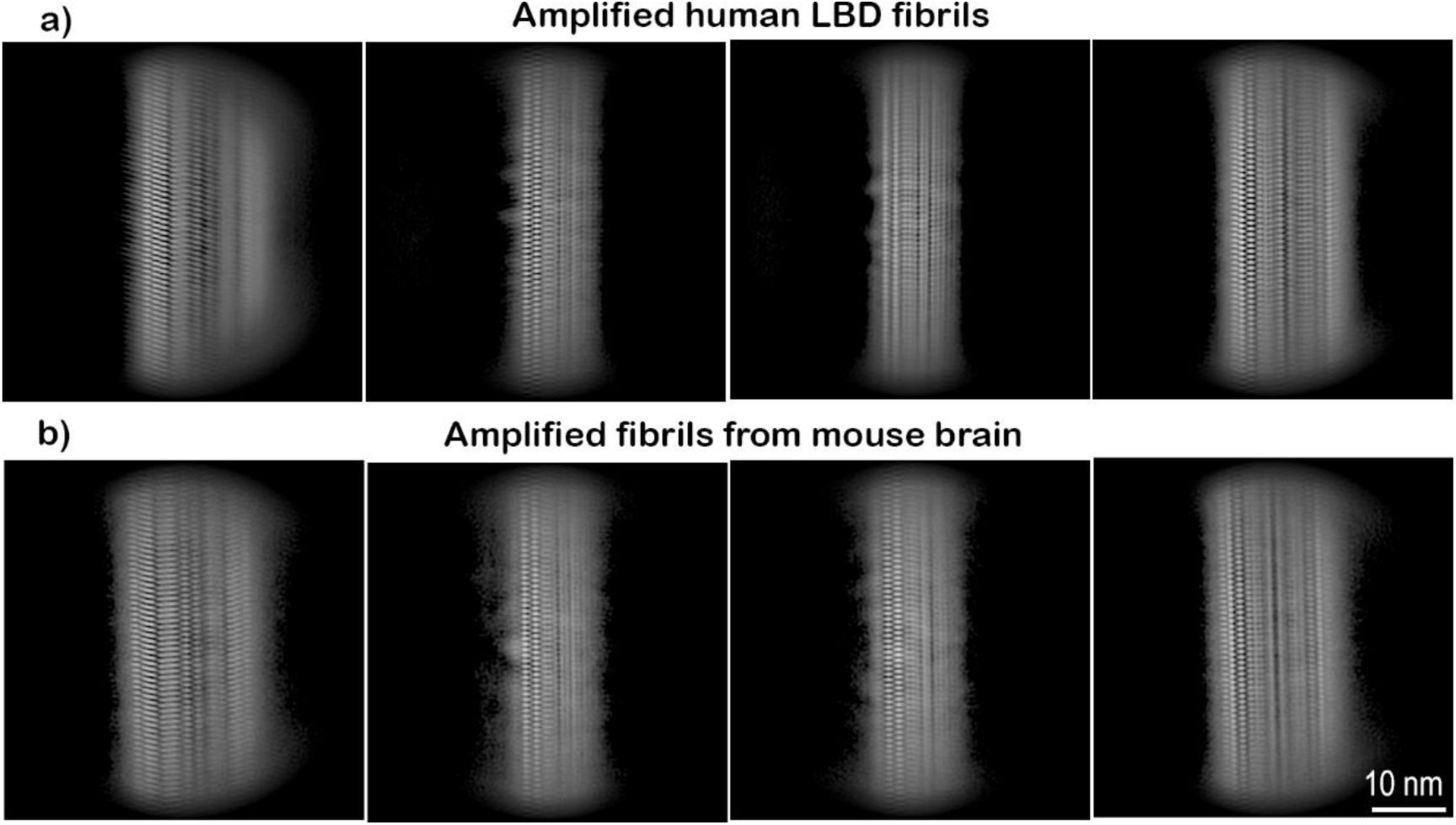
2D class averages from single particle cryo-EM data obtained for Asyn fibrils amplified from mouse brain tissue are highly similar to those obtained for amplified human LBD Asyn fibrils. Representative 2D class averages obtained for human amplified LBD fibrils from the LBD 1 case are shown in the top row (a) 2D class averages for amplified fibrils prepared from mouse brain tissue at 5 mpi following intrastriatal injection of human amplified LBD fibrils prepared from the LBD1 case are shown in the bottom row (b). The observed similarity between the two samples suggests that the protofilament fold for fibrils accumulating in mouse brain is highly similar to that of the amplified LBD fibrils from human postmortem brain tissue. Scale bar 10 nm.

## Discussion

We find that intrastriatal injection of an amplified LBD fibril preparation in a humanized Asyn mouse line results in progressive accumulation of Asyn fibrils that are structurally similar to the Asyn fibrils that accumulate in PD/LBD brain tissue. We also demonstrate that progressive accumulation of LBD Asyn fibrils is accompanied by neurodegeneration of substantia nigra dopaminergic neurons and motor impairment as measured by the rotarod test. These results establish a mouse model approach that enables further studies focused on mechanisms of fibril growth. The results also enable further studies of mechanisms by which LBD Asyn fibril accumulation leads to impaired neuronal function and neurodegeneration. Finally, this mouse model approach may be valuable in preclinical studies examining diagnostic and therapeutic approaches, a subset of which may be specific to fibril structure.

The development of mouse models based on injection of Asyn fibrils has advanced the study of disease mechanisms as well as diagnostic and therapeutic approaches in PD. The fibril injection approach typically provides consistent temporal and spatial patterns of fibril accumulation that can be utilized to study the effects of genetic and environmental manipulations on fibril accumulation[7,11,38,39]. The initial approach involved injecting an *in vitro* preparation of mouse Asyn fibrils into WT mice [7,10,13], with subsequent approaches using WT and mutant human Asyn fibril preparations injected into mouse strains with transgenic expression of WT or mutant human Asyn. Here we utilized an amplified LBD fibril preparation that has been shown to closely recapitulate the structure of LBD Asyn fibril isolated from postmortem LBD tissue. The observation that injection of amplified LBD fibrils results in a temporal and spatial pattern of fibril accumulation similar to the patterns observed after injection of *in vitro* Asyn fibril preparations in other mouse lines suggests that LBD fibrils and *in vitro* fibrils share some pathways for neuronal uptake, intracellular transport, fibril growth, and clearance mechanisms. Future studies using this humanized Asyn mouse line or other mouse lines can compare LBD Asyn fibrils with *in vitro* fibril preparations as well as MSA fibrils to further examine the degree to which patterns differ when compared in the same mouse line.

Comparison of 2D class averages for fibrils amplified from mouse brain tissue and the fibrils amplified from human LBD tissue used for injection indicates that the Lewy fold is maintained during fibril propagation in mouse neurons. Further analysis based on comparison of cryo-EM 3D density maps would provide further information on the degree of structural similarity between the two fibril preparations, which is currently a substantial challenge given their lack of helical twist. The observed structural similarity indicates that the intracellular environment in mouse neurons is permissive for maintaining the Lewy fold during 5 months of fibril accumulation after injections. The initial formation of fibrils with PD/LBD structure may depend on factors specific to the intracellular environment in PD neurons, as indicated by the fact that the Lewy fold has not been observed in Asyn fibrils prepared *in vitro* under a variety of condition[40]. However, environmental factors may not be required to maintain PD/LBD structure during fibril growth. The presence of nonproteinaceous densities in the cryo-EM structures of LBD fibrils isolated from postmortem PD/LBD brain tissue [37] suggests that fibril formation and growth could be modulated by molecules that associate with the fibrils. Alternatively, fibril growth may not require cofactors, and the observed association of nonproteinaceous molecules may occur ndependent of the growth process.

The mouse model approach described here can be useful in future studies of disease mechanisms and therapeutic approaches targeting Asyn fibril accumulation and its downstream consequences. While approaches based on *in vitro* fibril preparations have been and will continue to be useful for these studies, an approach that maintains LBD fibril structure may yield more informative results for structure-specific mechanisms and therapeutics. Further studies are needed to understand how often results may be affected by fibril structure, but its potential relevance is supported by previous studies [41,42]. Uemura et al [19] showed that injection of an *in vitro* PFF Asyn fibril preparation and an amplified LB Asyn fibril preparation produced distinct temporal and anatomic patterns of fibril accumulation in a Thy1: SNCA/Snca–/– mouse line overexpressing human Asyn under the Thy1 promoter. In addition, previous studies of Positron Emission Tomography (PET) ligand development for Asyn fibril accumulation demonstrate structure-dependent differences in the binding of small drug-like molecules to Asyn, which may also be relevant to small-molecule therapeutics targeting Asyn fibrils. For example, the radioligand Tg-1-90B binds with higher affinity to amplified LBD fibrils compared to an *in vitro* Asyn fibril preparation [43]. In addition, the [11C] M503-1619 radioligand bound to LBs in PD brain more robustly than GCIs in MSA brain [44], while the [11C] HY-2-15 radioligand showed specific binding of GCIs in MSA but low specific binding to LBs in PD/LBD brain tissues [45].

This study has several limitations. The structural characterization of Asyn fibrils accumulating in mouse brain tissue is limited to analysis of 2D class averages, which lacks the level of structural detail provided by either 3D cryo-EM reconstruction resolved to atomic resolution or SSNMR-derived structural models. The lack of twist for LBD Asyn fibrils limits 3D processing of the cryo-EM data. Resolution may be improved through further advances in cryo-EM data processing for non-twisting fibrils. Another potential limitation for future applications of this mouse model is the degree to which mice injected with LBD Asyn fibrils develop neurodegeneration and neurologic impairment. We observed significant deficits in rotarod performance and significant loss of dopaminergic neurons, but the effects were relatively modest and may not provide optimal sensitivity for testing therapeutic approaches that target fibril accumulation or downstream consequences of fibril accumulation. Alternative approaches utilizing injection of amplified LBD fibrils in mice with transgenic overexpression of human Asyn may produce more robust neurodegeneration resulting from LBD fibril accumulation.

Future studies can examine whether additional neurodegenerative changes are present in this mouse model. Cryo-ET studies that investigate Asyn fibril structure in situ can provide further information on the structure of accumulating fibrils and their subcellular location, as well as their association with organelles and proteins involved in clearance of fibrils. These studies can include short intervals after fibril injections to capture uptake and transport in neurons as well as non-neuronal cells, including microglia.

Future studies can also include injection of LBD Asyn fibrils in mouse strains overexpressing human Asyn, to determine whether LBD fibril structure is maintained in accumulating fibrils, and whether mice develop higher rates of fibril accumulation and greater levels of neurologic impairment.

## Acknowledgments

We thank the WashU Institutional Animal Care and Use Committee (IACUC) and the Department of Comparative Medicine (DCM) for their oversight, guidance, and support in ensuring the ethical and compliant conduct of this research. We thank the Mouse Genetics Core, Department of Pediatrics, for breeding the mice. We thank the Animal Behavior Core for the behavioral studies, with support from the McDonnell Center for Systems Neuroscience, McDonnell Center for Cellular and Molecular Neuroscience, and the Taylor Family Institute at Washington University in St. Louis. Immunostaining and cryo-EM imaging were performed in part using the Washington University Center for Cellular Imaging (WUCCI)

## Author Contributions

Conceptualization: Paul T. Kotzbauer, Rebecca Miller, Aditi V. Bagade, Dhruva D. Dhavale; Methodology: Paul T. Kotzbauer, Rebecca Miller, Aditi V. Bagade, Dhruva D. Dhavale; Formal analysis and investigation: Rebecca Miller, Dhruva D. Dhavale, Aditi Bagade, Joshua T. Dearborn, Katerine Basore, Summers Brock, Bradley Rednour, Jennifer O’Shea, Matthew Koelling; Writing - original draft preparation: Aditi Bagade, Paul T. Kotzbauer, Dhruva D. Dhavale; Writing - review and editing: all authors; Funding acquisition: Elizabeth Wright, Paul T. Kotzbauer; Resources: Paul T. Kotzbauer, Dhruva D. Dhavale; Supervision: Paul T. Kotzbauer, Rebecca Miller, Dhruva D. Dhavale.

## Data Availability

The data generated during this study are included in this article and its supplementary information. Additional data are available from the corresponding author upon request.

## Ethical Declaration

All animal procedures were performed in accordance with NIH guidelines [19] approved by the Institutional Animal Care and Use Committee (IACUC) at Washington University School of Medicine under the protocols 19-1098 and 23-0003. The human post-mortem tissue was clinically and neuropathologically characterized by the Movement Disorders Brain Bank, Washington University, St. Louis, MO. Written informed consent was obtained from all the participants to perform a brain autopsy. Confirmed consent for research purposes was obtained after death from the immediate next-of-kin for brain removal and retention of brain tissue.

