## Supplemental Data for "Intrastriatal injection of amplified Lewy body dementia (LBD) fibrils in mice results in accumulation of fibrils with LBD structure and neurodegeneration"

Supplementary figures:

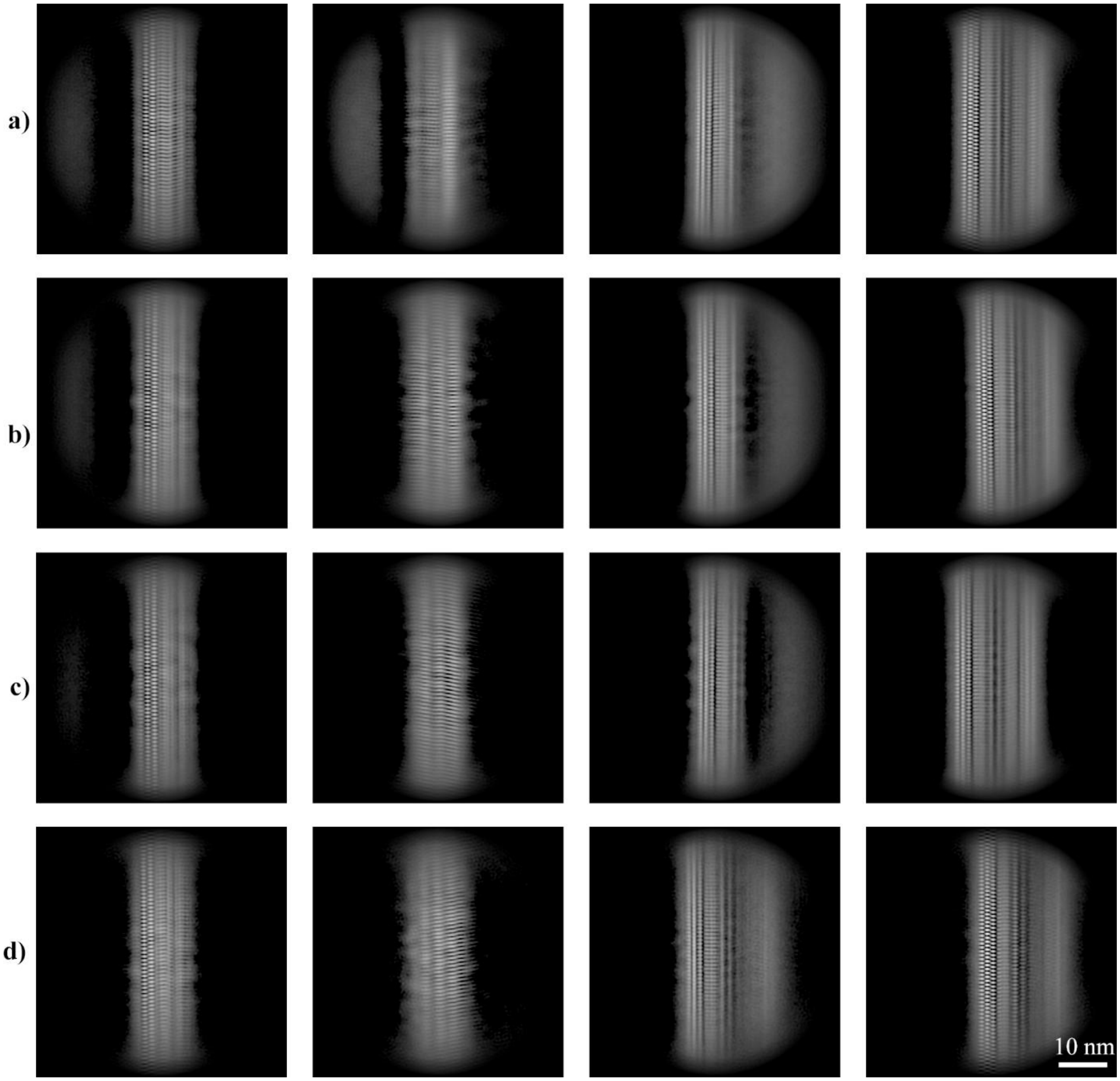


**Figure S1**: Cryo-EM 2D class averages of amplified LBD Asyn fibrils derived from fibril seeds prepared with different extraction methods show similar projections. The top row shows four 2D class averages derived from amplified LBD fibrils prepared with the insoluble fraction from the LBD1 case that was previously used for SSNMR structure studies (a). The second through fourth rows (b-d) show 2D class averages of amplified LBD Asyn fibrils prepared with sarkosyl-insoluble fractions from the LBD1, LBD7, and LBD8 cases, respectively. The highly similar projections between all samples are consistent with a shared protofilament fold. Scale bar 10 nm.

a)

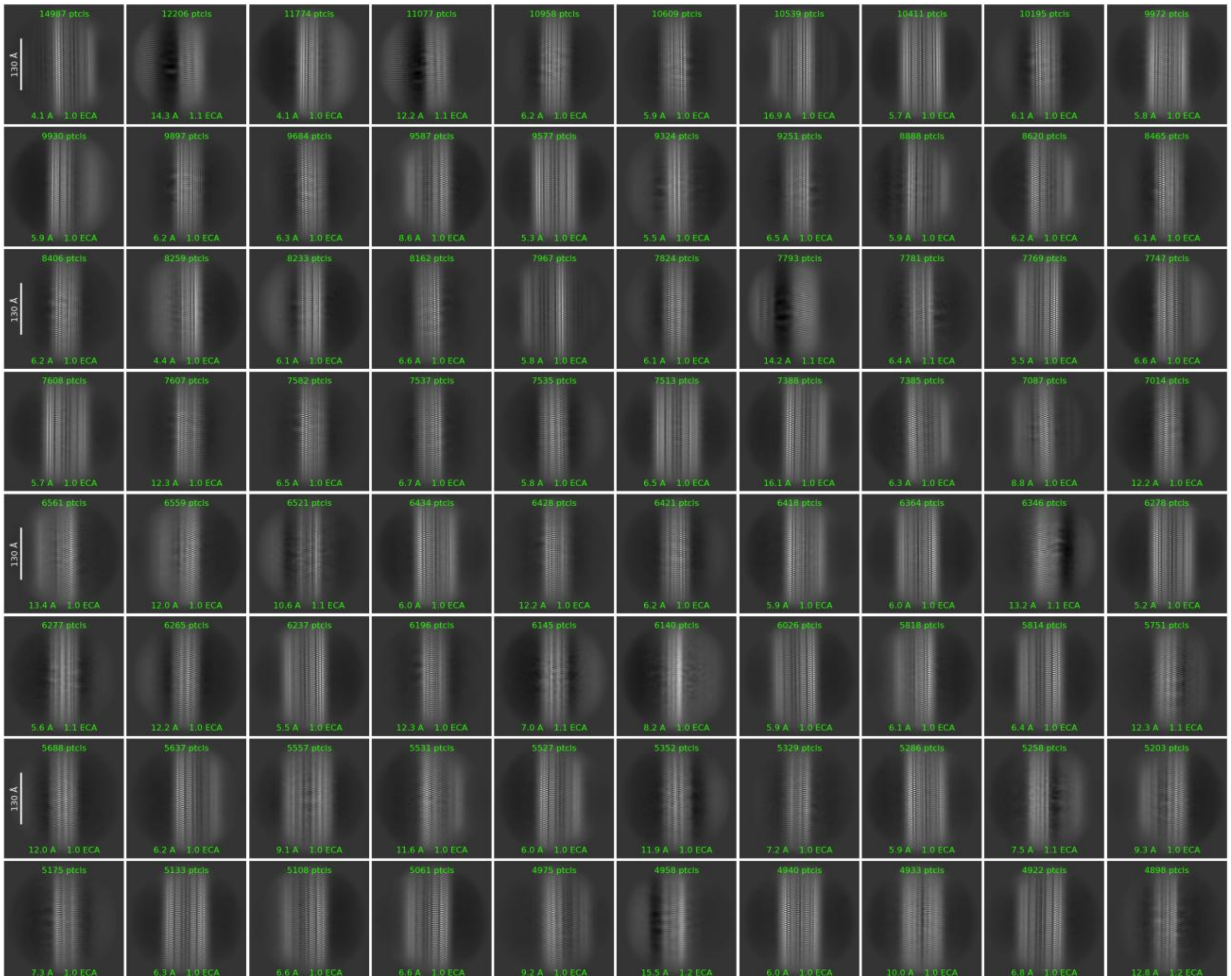

**b)**

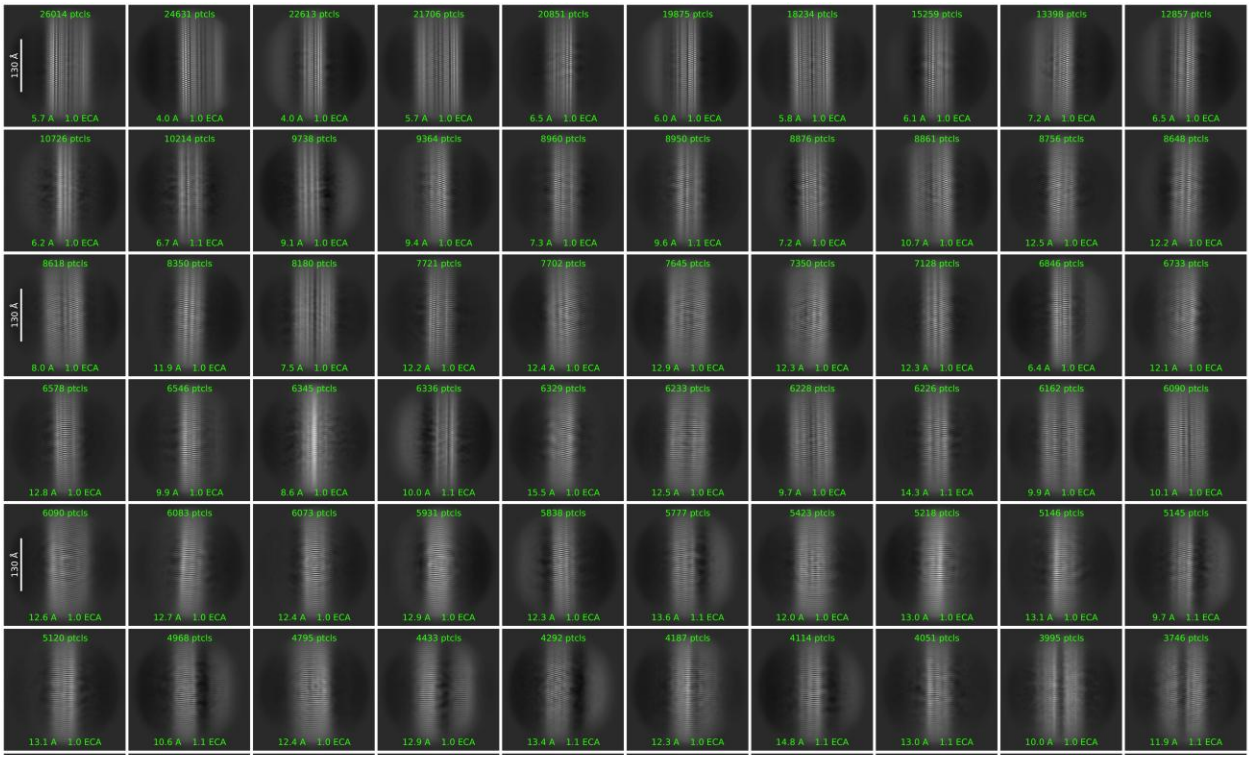

**c)**

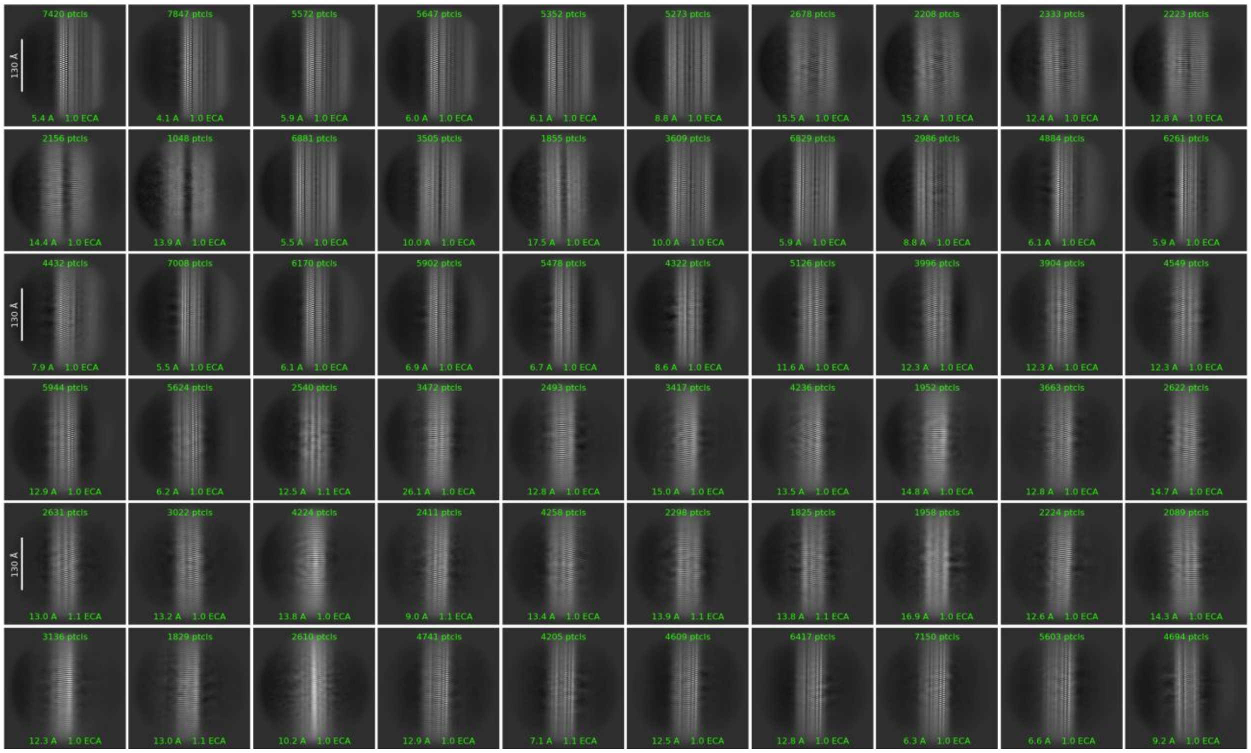

**d)**

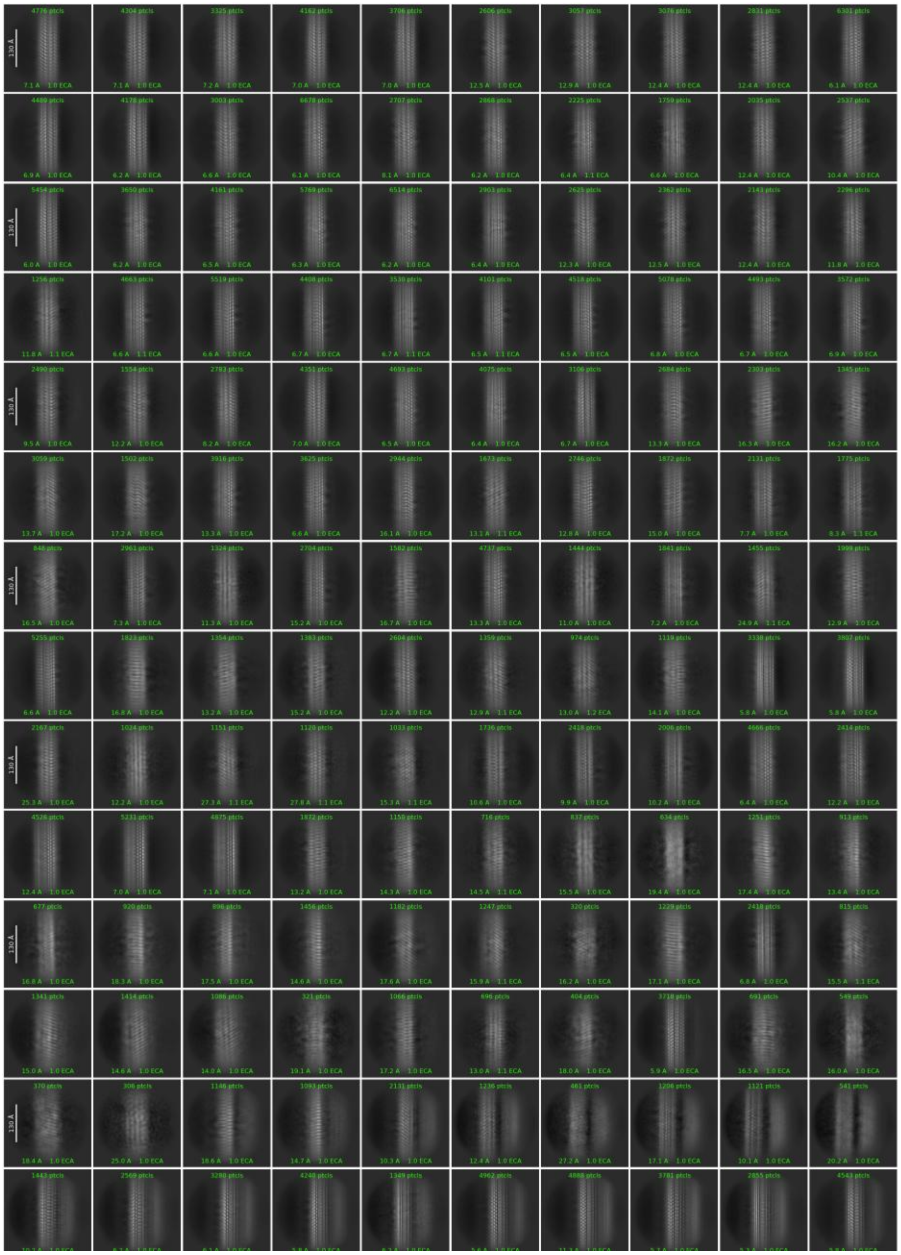

**Figure S2**: **2D class averages for human amplified LBD fibrils**. 2D classification of single-particle cryo-EM data was performed with cryoSPARC software and are arranged in order of the highest number of particles. (a) The 2D class averages derived from amplified LBD fibrils prepared with the insoluble fraction from the LBD1 case. The images used in Fig. 5 were obtained from positions (row/column) (B9, D3, B2, G8), and for Fig. S1 (E6*, H6, A3, C9*). (b-d) The 2D class averages of amplified LBD Asyn fibrils prepared with sarkosyl-insoluble fractions from the cases (b) LBD1, (c) LBD7, and (d) LBD8. The images used in Fig. S1 were obtained from positions for (b) A8, D8, A3*, A2; (c) D2*, E2, C2, B3; (d) C2, H6, N9, N7. Asterisks (*) denote images rotated 180º around the vertical axis. Scale bars: = 130 Å.

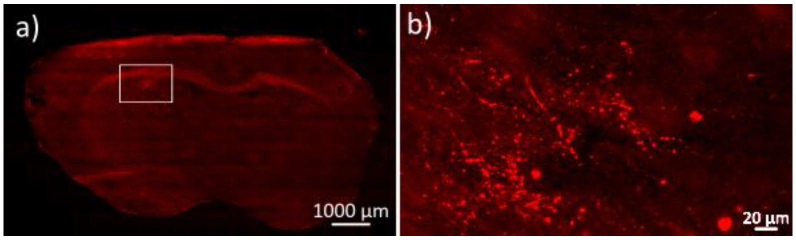


**Figure S3**: Immunostaining for pAsyn in mouse brain tissue sections at 7 days post-injection of amplified LBD fibrils. Representative coronal brain sections at 7 days following intrastriatal injection of amplified LBD fibrils, showing pAsyn aggregates near the injection site at (a) low magnification and (b) high magnification. The white box in (a) indicates the region magnified in (b). Scale bars: 1000 μm (a), 20 μm (b).

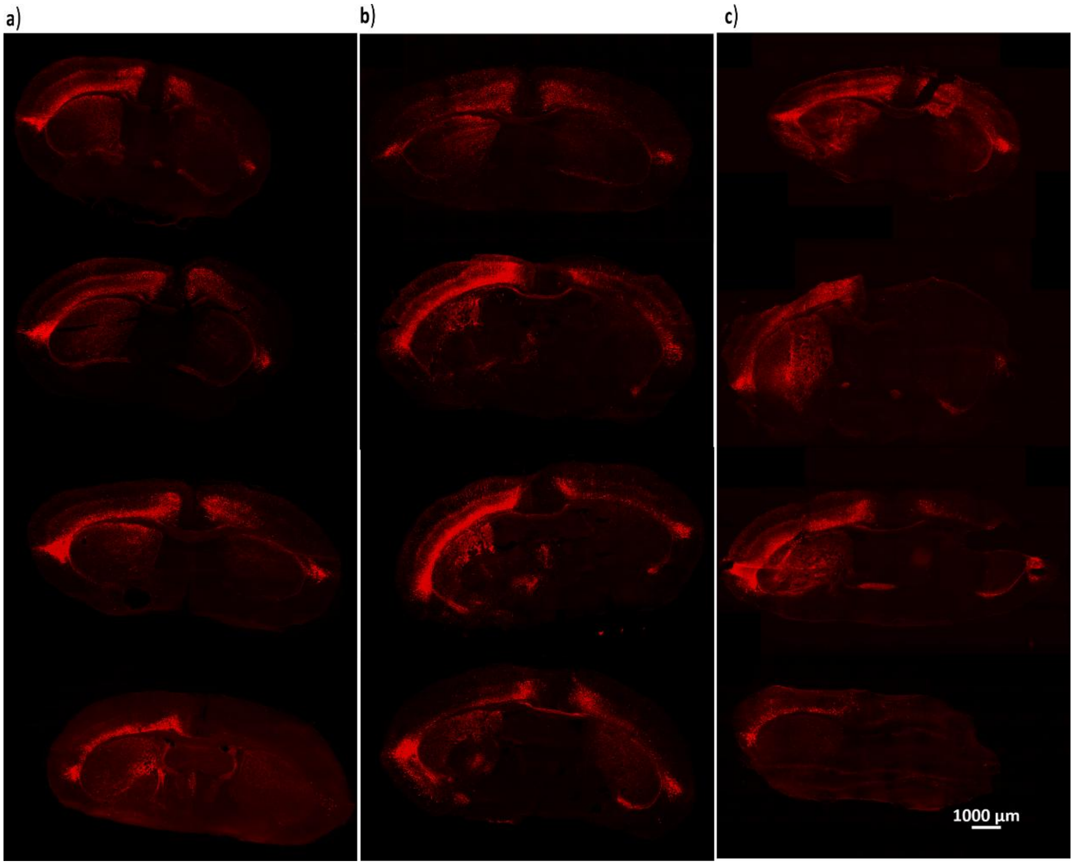


**Figure S4**: pAsyn immunostaining following intrastriatal injection of amplified LBD Asyn fibrils prepared from three postmortem LBD cases. Anti-pAsyn immunostaining results were observed for n=4 mice injected with amplified fibrils prepared from the LBD1 (a), LBD7 (b) and LBD8 (c) cases. Low-magnification images of coronal sections at Bregma are shown for each mouse 3 mpi. Scale bar: 1000 μm

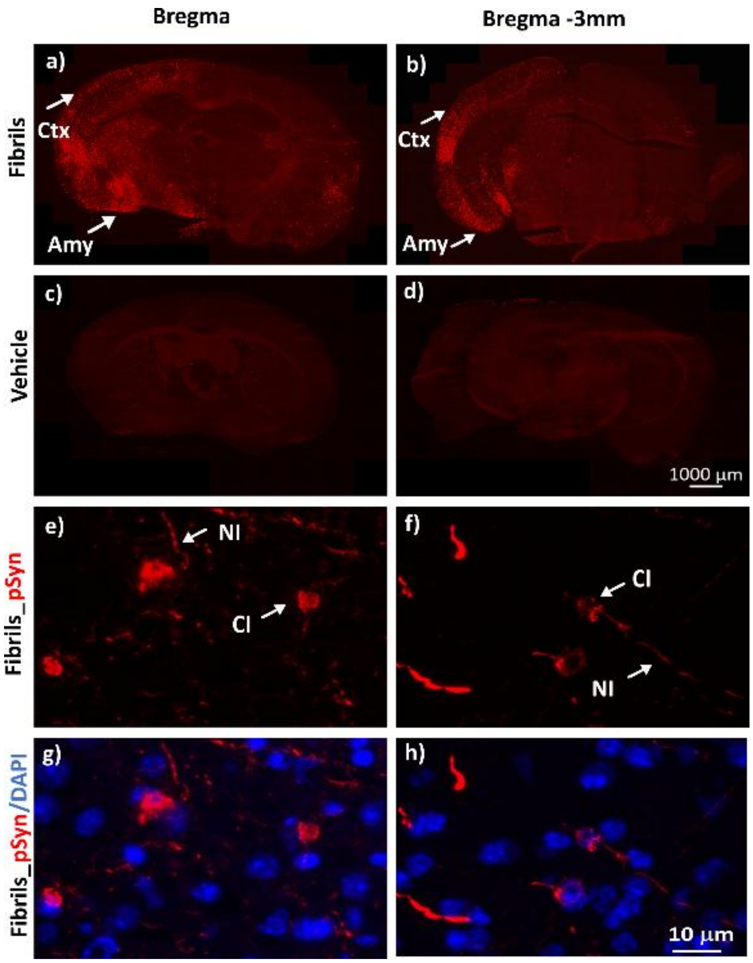


**Fig S5**: Persistence of Asyn fibril accumulation at 14 mpi after injection of amplified LBD fibrils, demonstrated by immunostaining for pAsyn (a, b). No pathological staining was observed in PBS-injected mice at 14 mpi (c, d). Mice injected with amplified LBD fibrils showed pAsyn cytoplasmic inclusions (CI) and neuritic inclusions (NI) in the cortex (e, f) following pAsyn immunostaining and DAPI labeling of nuclei (g, h). Note that CI appear more consolidated at 14 mpi compared to earlier time points (1-5 mpi) as shown in Fig. 1. Abbreviations: Cortex (Ctx), Striatum (Str), Amygdala (Amy); pAsyn (Red); DAPI (blue). Scale bars: 1000 µm (a-d), 10 µm (e-h).

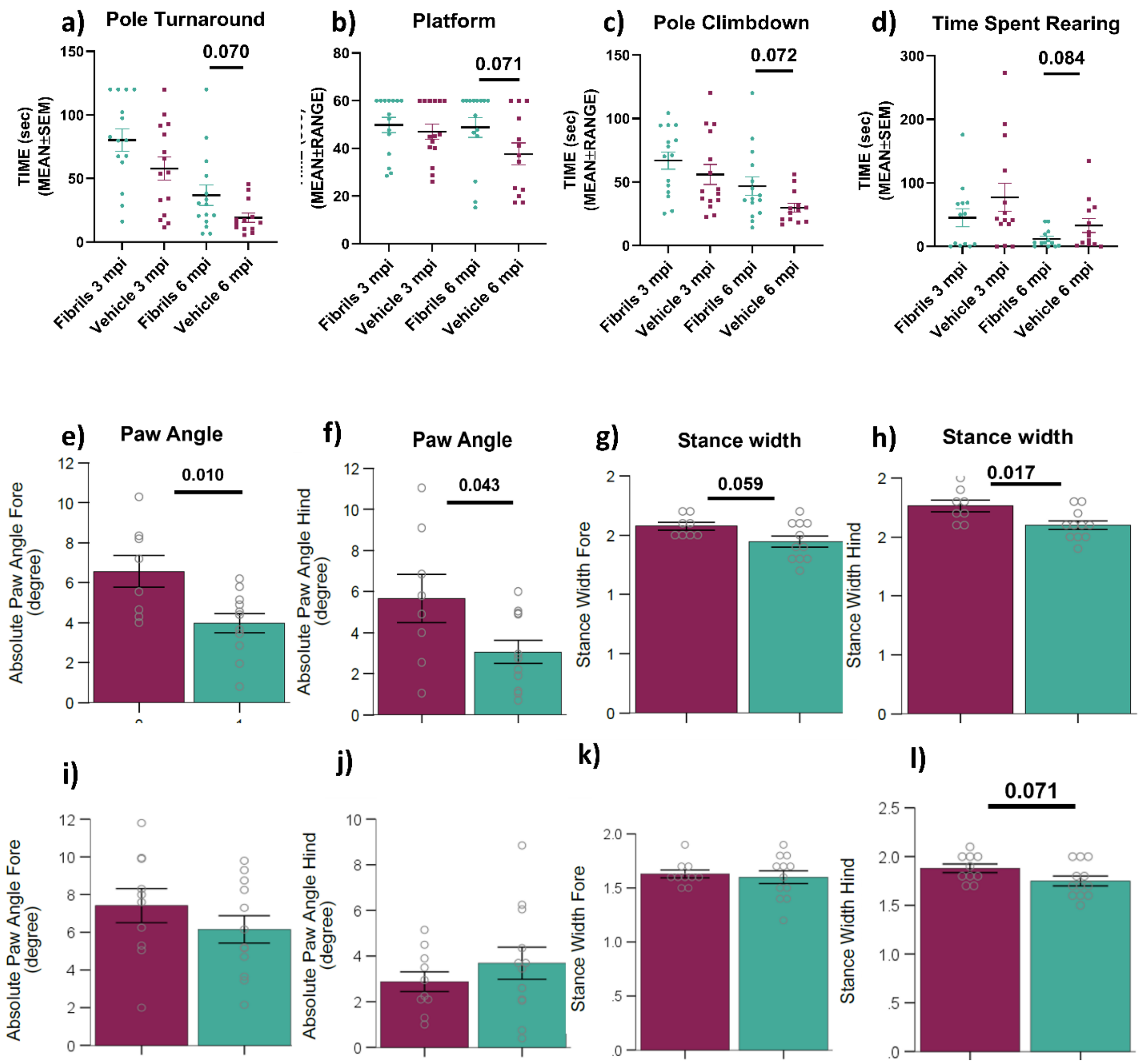


**Fig. S6**: Behavior analysis at 3 mpi and 6 mpi: (a-d) Locomotor measures did not show any significant differences but exhibited a trend of impaired performance by mice injected with fibrils at 6 mpi as compared to control mice injected with phosphate buffer saline. Gait analysis at 3 mpi (e-h) and 6 mpi (i-l) showed Significant differences in forepaw angle (e), hind paw angle (f), and hind paw stance width (h) at 3 mpi. Whereas no significant differences were observed at 6 mpi (i-l, and g). An unpaired two-tailed t-test was used for statistical analysis. Data are presented as mean ± SEM. Control-injected mice are shown in maroon, while fibril-injected mice are shown in green.

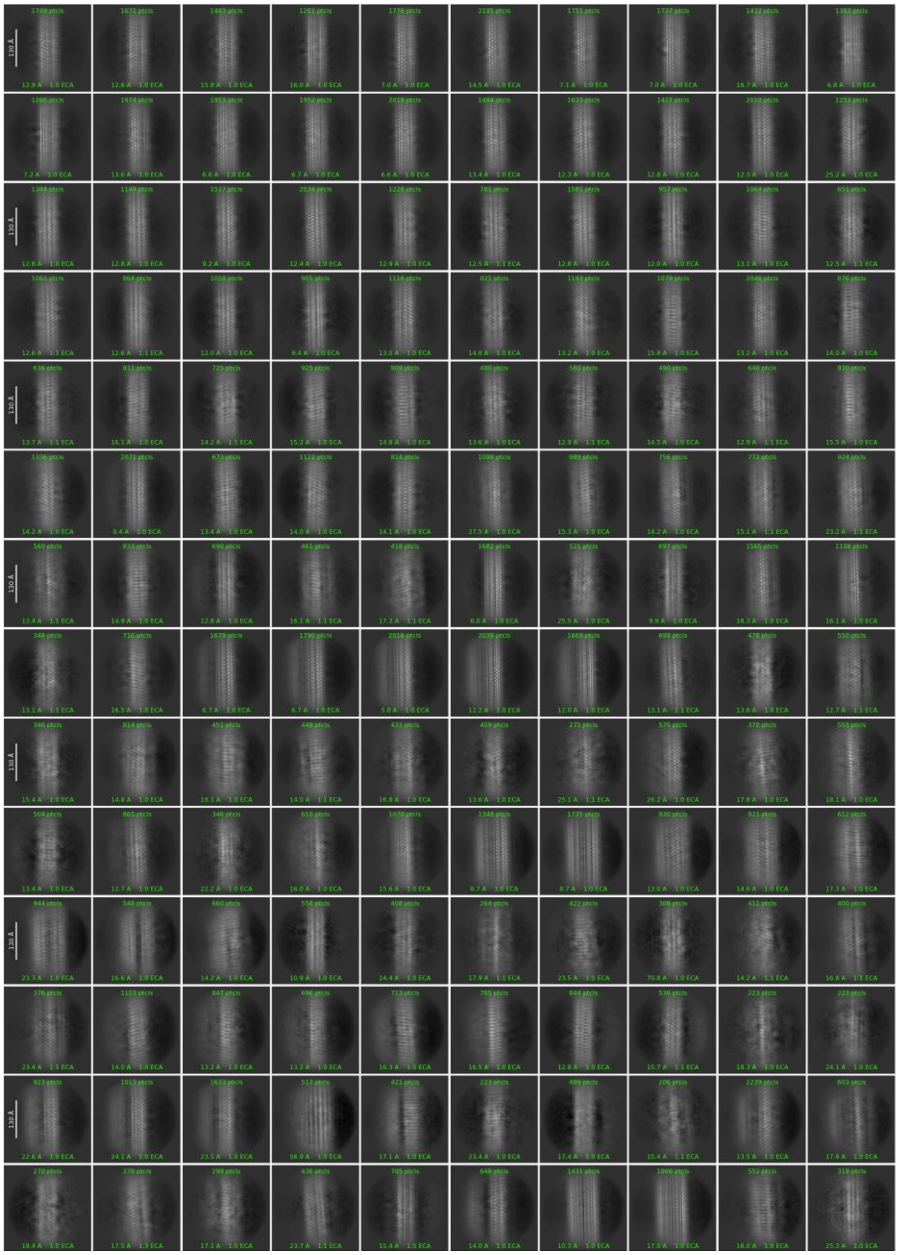

**Figure S7**: 2D class averages for fibrils amplified from mouse brain tissue following intrastriatal injection of amplified LBD fibrils. 2D classification of single particle cryo-EM data was performed with cryoSPARC software as outlined in the methods. 2D class averages for the one hundred forty 2D classes with the highest number of particles are shown. The images used in Fig. 5 were obtained from positions (row/column) A8, B4, J6*, and N8. Asterisks (*) denote images rotated 180º around the vertical axis. Scale bars 130 Å.

Supplementary Table 1: Demographics of human cases used in the study

| Case | Sex (Male /Female) | Age at Death, median (range) | Disease Duration (years) | Postmortem interval (hours) | Primary Diagnosis | LB Stage (0-6) | Amyloid Stage 0,(A-C) | BRAAK NFT Stage (I-VI) |
|---|---|---|---|---|---|---|---|---|
| LBD1 | M | 77 (71-82) | 6 | 14 | LBD | 6 | C | II |
| LBD7 | M | 77 (71-82) | 10 | 11 | LBD | 6 | C | III |
| LBD8 | M | 72 (71-82) | 10 | 5.5 | LBD | 5 | 0 | I |

Supplementary Table 2: Antibodies used in immunohistochemistry studies.

| Primary/Secondary Antibodies | Source/Cat No | Host and clonality | Antibody conc. | Incubation time |
|---|---|---|---|---|
| Biotin anti-alpha synuclein phosphor Ser 129 antibody | Biolegend/825704 | Mouse Monoclonal | 1.25 µg/mL | Overnight at 4ºC |
| Anti-Tyrosine hydroxylase antibody | Millipore Sigma/AB152 | Rabbit Polyclonal | 0.5 µg/mL | Overnight at 4ºC |
| Alexa Fluor 594 Streptavidin | Biolegend/405240 | Not applicable | 1µg/mL | 1h at RT |
| Donkey Anti-rabbit (Alexa Fluor 488) | Invitrogen/A21206 | Donkey Polyclonal | 4µg/mL | 1h at RT |
